# Comparative Analysis of *Drosophila* Bam and Bgcn Sequences and Predicted Protein Structural Evolution

**DOI:** 10.1101/2024.12.17.628990

**Authors:** Luke R Arnce, Jaclyn E Bubnell, Charles F Aquadro

## Abstract

The protein encoded by the *Drosophila melanogaster* gene *bag of marbles* (*bam*) plays an essential role in early gametogenesis by complexing with the gene product of *benign gonial cell neoplasm* (*bgcn*) to promote germline stem cell daughter differentiation in males and females. Here, we compared the AlphaFold2 and AlphaFold Multimer predicted structures of Bam protein and the Bam:Bgcn protein complex between *D. melanogaster, D. simulans, and D. yakuba*, where *bam* is necessary in gametogenesis to that in *D. teissieri*, where it is not. Despite significant sequence divergence, we find very little evidence of significant structural differences in high confidence regions of the structures across the four species. This suggests that Bam structure is unlikely to be a direct cause of its functional differences between species and that Bam may simply not be integrated in an essential manner for GSC differentiation in *D. teissieri*. Patterns of positive selection and significant amino acid diversification across species is consistent with the Selection, Pleiotropy, and Compensation (SPC) model, where detected selection at *bam* is consistent with adaptive change in one major trait followed by positively selected compensatory changes for pleiotropic effects (in this case perhaps preserving structure). In the case of *bam*, we suggest that the major trait could be genetic interaction with the endosymbiotic bacteria *Wolbachia pipientis*. Following up on detected signals of positive selection and comparative structural analysis could provide insight into the distribution of a primary adaptive change versus compensatory changes following a primary change.

## INTRODUCTION

Gamete production in most sexually producing animals starts with germline stem cell (GSC) division, in which GSCs divide asymmetrically producing one daughter to self-renew, maintaining the germline, and another to differentiate into sperm and eggs (Kahney et al. 2019). Effective regulation of these GSC divisions is fundamental for proper reproduction, and improper regulation of this process can lead to sterility (Gleason et al. 2018). Consequently, GSC divisions are highly regulated and presumed to be highly conserved. However, several genes essential for reproduction whose products control GSC maintenance and differentiation in *D. melanogaster* are rapidly evolving due to positive selection for amino acid diversification (Civetta et al. 2006; Bauer DuMont et al. 2007; Choi and Aquadro 2015; Flores, DuMont, et al. 2015). Some of these genes whose function is essential for reproduction in the *D. melanogaster* germline are nonessential for fertility, or even absent, in other *Drosophila* species (Bubnell et al. 2022).

One key GSC gene is *bag-of-marbles* (*bam*). *Bam* encodes a 442 amino acid protein that is predicted to be partially disordered and functions in hematopoiesis, maintenance of gut integrity, and as the key switch gene in *D. melanogaster* for GSC daughter differentiation in females and for spermatogonia terminal differentiation in males (McKearin and Spradling 1990; Civetta et al. 2006; Bauer DuMont et al. 2007; Insco et al. 2009; Tokusumi et al. 2011).

In *D. melanogaster* female GSCs, *bam* is initially repressed by extrinsic DPP signaling in the stem cell niche. Upon GSC division, movement of the daughter cell away from the cap cells enables transcription of *bam* (Chen et al. 2003) and differentiation into a cystoblast. Bam binds to the protein product of *benign gonial cell neoplasm* (*bgcn*) in cystoblasts to promote differentiation (Li et al. 2009). Bam’s binding with other protein partners represses the translation of self-renewal factors *nanos* and *eIF4a* (Li et al. 2009; Chau et al. 2012; Xie 2013; Malik et al. 2020). Cystoblasts then undergo four mitotic divisions with Bam concentrated in the fusome, a structure connecting the sister cells composing the cyst. Bam and Bgcn function together to regulate mitotic synchrony in these cells (Chen and McKearin 2003). Prior to meiosis, *bam* expression is repressed.

During spermatogenesis in *D. melanogaster* males, the spatiotemporal expression of *bam* and its subsequent function in differentiation slightly differs from its role in oogenesis. *Bam* is expressed in GSCs and, as differentiation progresses, expression of *bam* increases (Insco et al. 2009). Once Bam protein reaches a threshold in dividing spermatogonia, its binding with Bgcn represses *mei-P26*, triggering the switch to terminal differentiation, followed by meiosis (Chen et al. 2014) *Bam* expression is then quickly repressed.

In both *D. melanogaster* males and females, Bam’s interaction with CAF40, a part of the CCR4-NOT deadenlyase complex, could contribute to *nanos* mRNA destabilization (Fu et al. 2015; Sgromo et al. 2017). This inhibition of *nanos* translation promotes GSC differentiation. Self-renewal is also repressed through Bam’s interaction with Csn4 (Pan et al. 2014). The COP9 signalosome regulates many signaling and cell cycle pathways in Drosophila and is essential for regulating gene expression during development as well as having essential roles in oogenesis and ovarian germline stem cells in *Drosophila melanogaster*. It consists of nine Csn subunits in GSCs. Bam expression in differentiating daughter cells sequesters subunit Csn4 from the COP9 complex, inhibiting the complex’s function in GSC self-renewal (Freilich et al. 1999).

Additionally, Bam forms a complex with Otu to promote GSC differentiation by deubiquitinating Cyclin A (CycA) (Ji et al. 2017). This stabilizes CycA levels and promotes differentiation of GSC daughter cells. Loss of *bam* function in both male and female *D. melanogaster* results in overproliferation of spermatogonia or GSCs in males and females, respectively, leading to severe fertility defects causing sterility (McKearin and Spradling 1990).

Despite *bam*’s essential role in GSC daughter differentiation in *D. melanogaster*, recent results indicate the gene is not highly conserved in sequence or function even among closely related species (Figure 1). Between sibling species *D. melanogaster* and *D. simulans*, there are 60 fixed amino acid differences out of the total 442 amino acids of Bam protein. Lineage-specific statistical tests of selective neutral molecular evolution indicate that 94% and 72% of these amino acid fixations were driven to fixation by natural selection between *D. melanogaster* and *D. simulans* and their common ancestor (Bubnell et al. 2022). Similar strong signals of positive natural selection are seen in other diverse lineages including those leading to *D. yakuba, D. serrata, D. birchi, D. jambulina, D. ananassae, D. americana* and *D. rubida* yet other lineages across the genus show no evidence of positive selection for protein diversification (Fig. 1 and Bubnell et al 2022). More distantly related *Drosophila* species differ from *D. melanogaster* by as much as 67% of their Bam amino acid sequence, and two outgroup species in which orthologs of Bam can be identified (*L. cuprina* and *M. domestica*) differ at 85% and 87% of their Bam protein sequence (Fig. 1).

**Figure 1.**
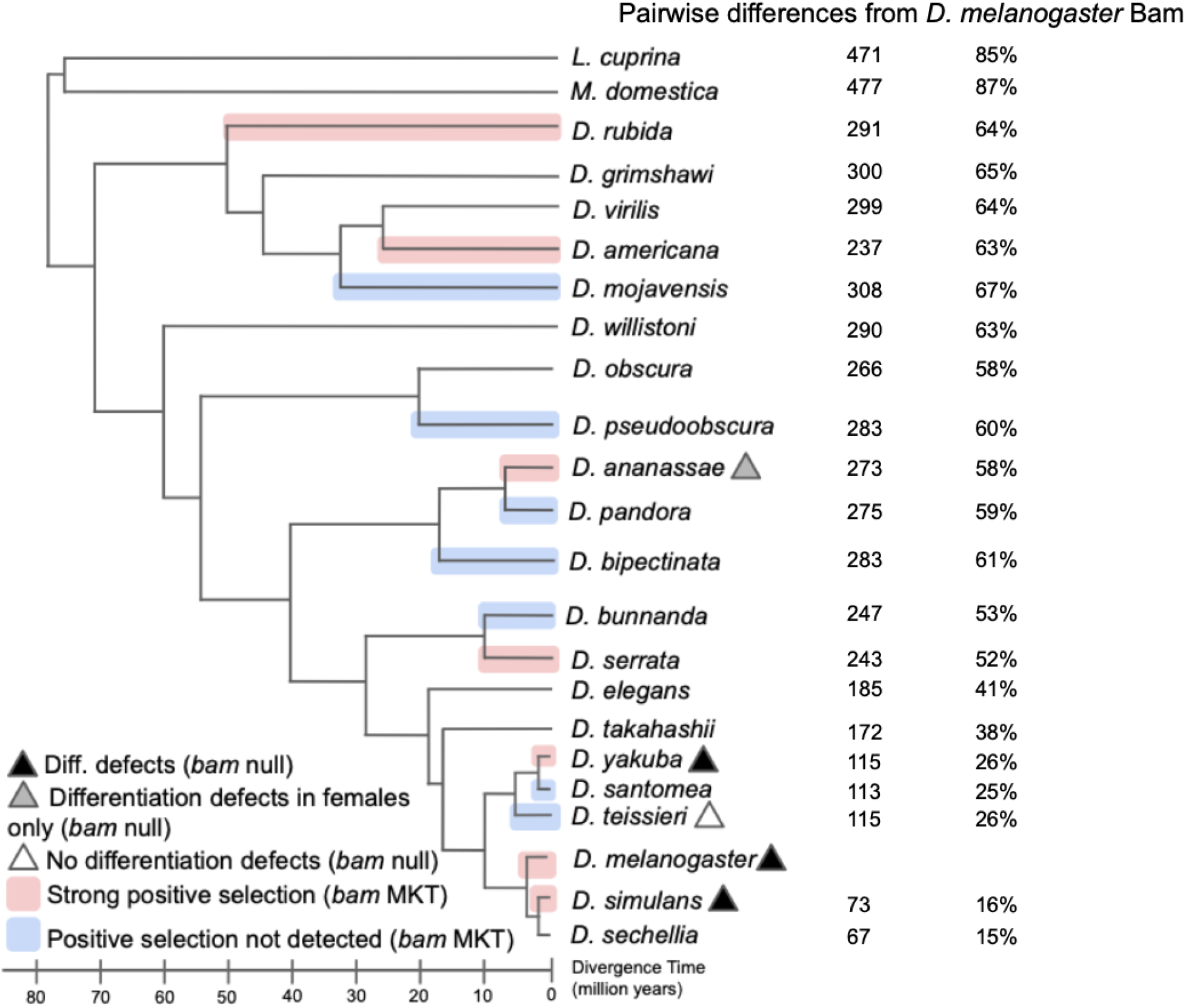
Heterogenous signals of positive selection and function in germline differentiation at *bam* inferred by the McDonald-Kreitman Test (MKT) and phenotypic assessment of *bam* null alleles across the *Drosophila* genus (Bubnell et al. 2022). Functional analysis of *bam* null mutants shows a range of differentiation defects with defects in *D. simulans, D. melanogaster*, and *D. yakuba*, differentiation defects in females only in *D. ananassae*, and no differentiation defects in both males and females in *D. teissieri*, demonstrating functional differences for *bam* across species. Pairwise amino acid differences from *D. melanogaster* Bam to other *Drosophila* and outgroup species show divergence ranging from 15% in the closely related *D. sechellia*, to 87% in the outgroup species *M. domestica*. Note that in pairwise alignments to outgroup species, pairwise differences are able to exceed 442 residues because the Bam protein in these species is comparatively larger than in *D. melanogaster*.

Recent generation of and functional analysis of *bam* null alleles in multiple species (Bubnell et al. 2022) demonstrated divergence in *bam* function among *D. simulans, D. melanogaster, D. teissieri, D. yakuba*, and *D. ananassae*. No differentiation defects were detected in male or female *D. teissieri bam* null mutants, indicating that *bam* does not function as an essential key switch gene for GSC differentiation in this species. Similarly, in *D. ananassae, bam* null females were sterile, although *bam* null males had normal spermatogenesis in males. These data together indicate *bam* sequence and function are both changing despite *bam*’s essential GSC function in several *Drosophila* species (Bubnell et al. 2022).

Given the divergence in *bam* function and high number of amino acid differences among *Drosophila* species, we hypothesized that the protein structures of orthologous *bam* proteins may be significantly different among species. We were particularly interested to determine whether the structure of Bam in *D. teissieri* differed from that in *D*. melanogaster and two other species where it has an essential role in the germline. Bam’s structure has been difficult to evaluate experimentally due to large amounts of disorder within the protein which have prevented structural determination using X-ray crystallography except for a small segment of Bam’s N-terminus (Sgromo et al. 2018). However, recently developed artificial intelligence computational methods including AlphaFold2, AlphaFold Multimer, and ChimeraX enable us to make predictions regarding Bam’s 3D protein structure (Jumper et al. 2021; Evans et al. 2021; Mirdita et al. 2022; Meng et al. 2023). Here, we focused our analysis on four *Drosophila* species (*D. melanogaster, D. simulans, D. teissieri*, and *D. yakuba*) with divergence time of up to 10 MYA in which *bam* function has been directly evaluated and found to be divergent (Bubnell et al. 2022). This group of species enables comparison of predicted Bam structure between species in which *bam* is either essential or not necessary for fertility in males and females.

Bam performs its key function as a switch gene for GSC differentiation as a protein complex with Bgcn (Li et al. 2009). Bgcn is a large protein (1215 amino acids) and is more ordered than Bam, but no X-ray crystallography structural data for it are available. We thus also generated predicted protein structures for Bgcn alone and the protein Bam:Bgcn complex. Significant differences in their structures could potentially be related to differences in Bam’s essential GSC switch function.

Here, we report that despite the extensive adaptively driven protein sequence divergence among all four species and the absence of a role of Bam in GSC differentiation in *D. teissieri*, the predicted structures for Bam, Bgcn and the Bam:Bgcn complex appear to be largely conserved. It is possible that the strong signature of natural selection at *bam* is associated with both a primary focal point of natural selection, perhaps related to Bam’s functional interaction with the intracellular bacteria *Wolbachia* (Flores et al. 2015, Russell et al. 2023), and secondary natural selection for compensatory mutations to preserve optimal protein structure. Major alteration of protein structure also does not appear to account for the lack of Bam’s role in GSC differentiation and fertility in *D. teissieri*.

## MATERIALS AND METHODS

### Sequences and alignments

Bam and Bgcn amino acid sequences for *D. melanogaster, D. simulans, D. teissieri*, and *D. yakuba* were identified via Basic Local Alignment Search Tool (BLAST) and extracted from their NCBI reference genomes (Camacho et al. 2009, O’Leary et al. 2016). MUSCLE was used to generate Bam and Bgcn multiple sequence alignments among these four species as well as an extended alignment for *bam* including 21 *Drosophila* species and 2 outgroup species (Edgar et al. 2004) (S1, github.com/la424/bam_bgcn).

Bam ancestral sequences predicted by Bubnell et al. (2022) were used for three phylogenetic nodes of interest: node A including all four species, node B including *D. teissieri* and *D. yakuba*, and node C including *D. melanogaster* and *D. simulans*. Bgcn ancestral sequences were generated with codeml at the same three nodes of interest by maximum likelihood (Fig 1, 2a, S2 github.com/la424/bam_bgcn).

**Figure 2a.**
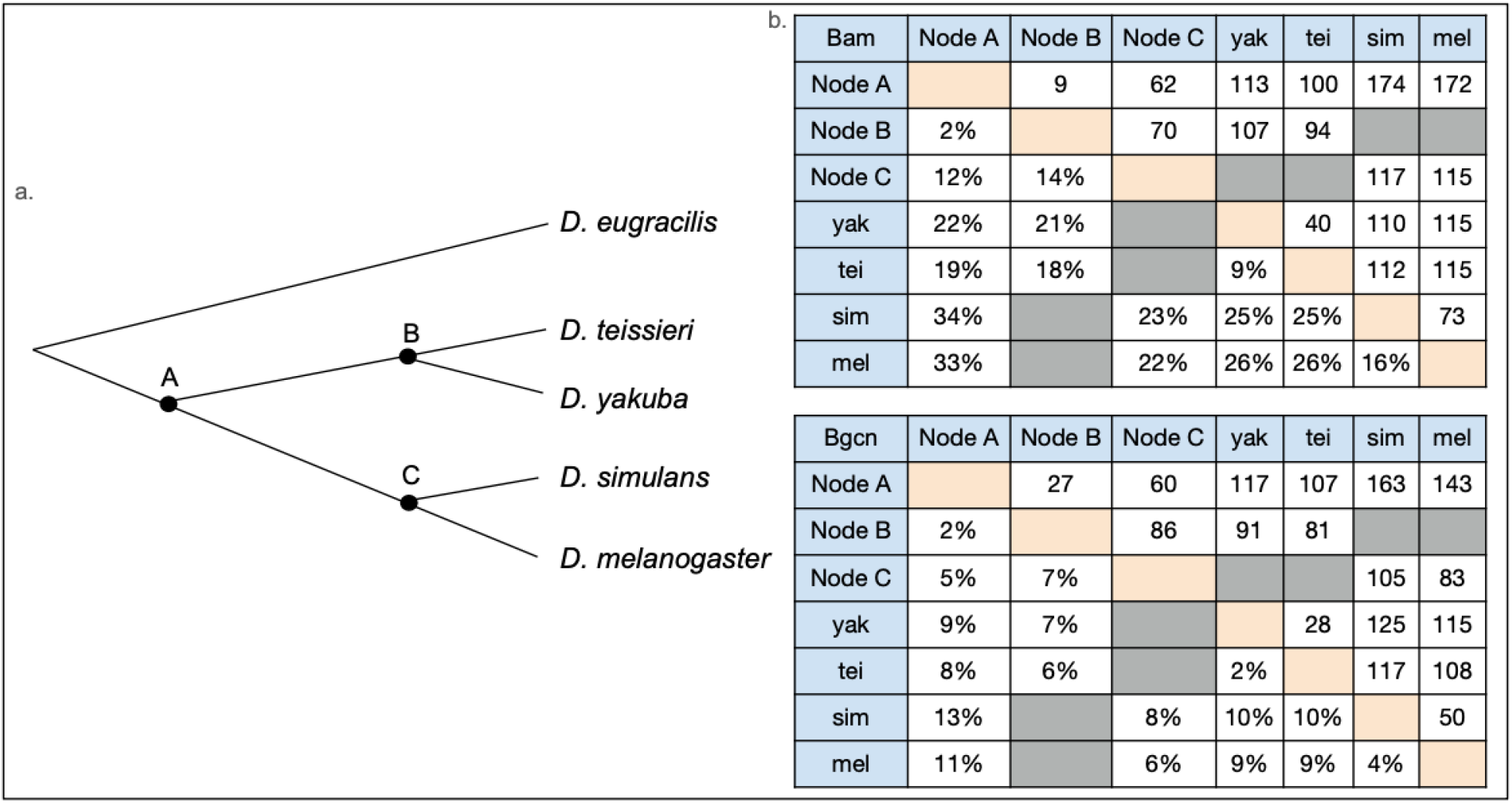
Cladogram of *Drosophila* species evaluated and outgroup (*D. eugracilis*) in this study and nodes with estimated ancestral amino acid sequences indicated as A, B, and C. b. Table of pairwise Bam and Bgcn amino acid sequence differences as raw numbers and percentages for *D. yakuba, D. teissieri, D. simulans, D. melanogaster*, and ancestral sequences A, B, and C.

### AlphaFold2 and AlphaFold multimer

AlphaFold predicted structures for Bam, Bgcn, and AlphaFold multimer predicted structures for Bam:Bgcn were generated with ColabFold v1.5.5:AlphaFold2 using MMseqs2 for extracted Bam and Bgcn amino acid sequences for the four included *Drosophila* species (Mirdita et al. 2022).

### ADOPT inference of structured vs disordered protein regions

ADOPT scores were generated for Bam and Bgcn amino acid sequences from *D. melanogaster, D*.*simulans, D. teissieri*, and *D. yakuba* using the ADOPT webserver (https://adopt.peptone.io/) (Accessed 16 December 2024) and resultant residue-specific values were graphed from sequence alignments using custom R scripts (S1, github.com/la424/bam_bgcn).

### Paired structural alignments between species

AlphaFold2 predicted structures were paired using Matchmaker in ChimeraX 1.7.1 with default settings (Meng et al. 2023). Paired alignments between each included species were generated for Bam, Bgcn, and Bam:Bgcn and assessed for differences using the metric root mean square deviation (RMSD) between paired residues in their current 3D positions.

Paired residues with less than 2 angstroms RMSD were classified as matched residues with no significant structural difference (Meng et al. 2023). Paired residues with more than 2 angstroms RMSD were classified as significantly different in structure. plDDT scores were incorporated to determine whether differences in paired structural alignments were in high confidence regions (github.com/la424/bam_bgcn).

### Predicted Hydrogen bonds within or between proteins

The H-bonds function in ChimeraX 1.7.1 was used with default functions to identify putative hydrogen bonding within Bam, Bgcn, and Bam:Bgcn for *D. melanogaster, D. simulans, D. teissieri*, and *D. yakuba*. Hydrogen bonds are called based on prediction metrics from high-resolution small-molecule crystal structures (Mills and Dean 1996). Distance tolerance (0.4 angstroms) and angle tolerances (20 degrees) for bonding were relaxed to identify hydrogen bonding within larger protein structures. H-bonds within protein structures were cataloged if bonded residues were more than five residues apart. H-bonds between complexed proteins, as in Bam:Bgcn, were all cataloged. H bond results for each of the predicted structures were then incorporated with the aligned sequences of the included species to enable characterization of differences in hydrogen bonding (github.com/la424/bam_bgcn, S4, S5).

Hydrogen bond differences were evaluated using several categories to search for potential patterns in significant differences. Hydrogen bonds were evaluated and compared by total number, by number in Bam and Bgcn functional and binding regions, by number in ordered and disordered regions, and by the ratio of bonds in ordered regions to disordered regions. Fisher’s exact tests followed by a Bonferroni correction for multiple hypothesis tests were then used to identify significant differences in hydrogen bonding for Bam, Bgcn, and Bam:Bgcn across species (github.com/la424/bam_bgcn, S4, S5).

### AlphaFold predicted contacts between Bam and Bgcn protein structures

The Alphafold Contacts function in ChimeraX 1.7.1 was used with default functions to identify residue pair interface points between Bam and Bgcn protein structures (Meng et al. 2023). The function uses PAE to determine confidence in relative position as well as distance in angstroms between the contact residues. Alphafold contacts with PAE values below 5 were compiled and incorporated with the aligned sequences of the included species to enable characterization of differences in AlphaFold contacts. AlphaFold contact differences were evaluated for Bam:Bgcn complexes across included species using the same strategy and categories outlined for evaluating significant differences in H-bonds (github.com/la424/bam_bgcn, S4, S5).

## RESULTS

### *Drosophila melanogaster* AlphaFold predictions for Bam, Bgcn, and the Bam:Bgcn complex

AlphaFold-predicted protein structures for Bam and Bgcn in *Drosophila melanogaster* reveal Bam as a partially disordered protein with ordered regions and Bgcn as a largely ordered, densely packed protein (Fig. 3). These structures and their interactions can be evaluated in the context of previous yeast 2-hybridization, co-immunoprecipitation, and GST pulldown to identify functional and interacting regions for the two proteins. (Li et al. 2009; Ji et al. 2009; Pan et al. 2014; Ji et al. 2017; Sgromo et al. 2017; Sgromo et al. 2018). Bam’s Bgcn binding regions were identified using pulldown assays (Pan et al. 2014). This direct experimental evidence of Bam and Bgcn interacting regions was used as a quality check for our Bam:Bgcn multimer predictions.

**Figure 3.**
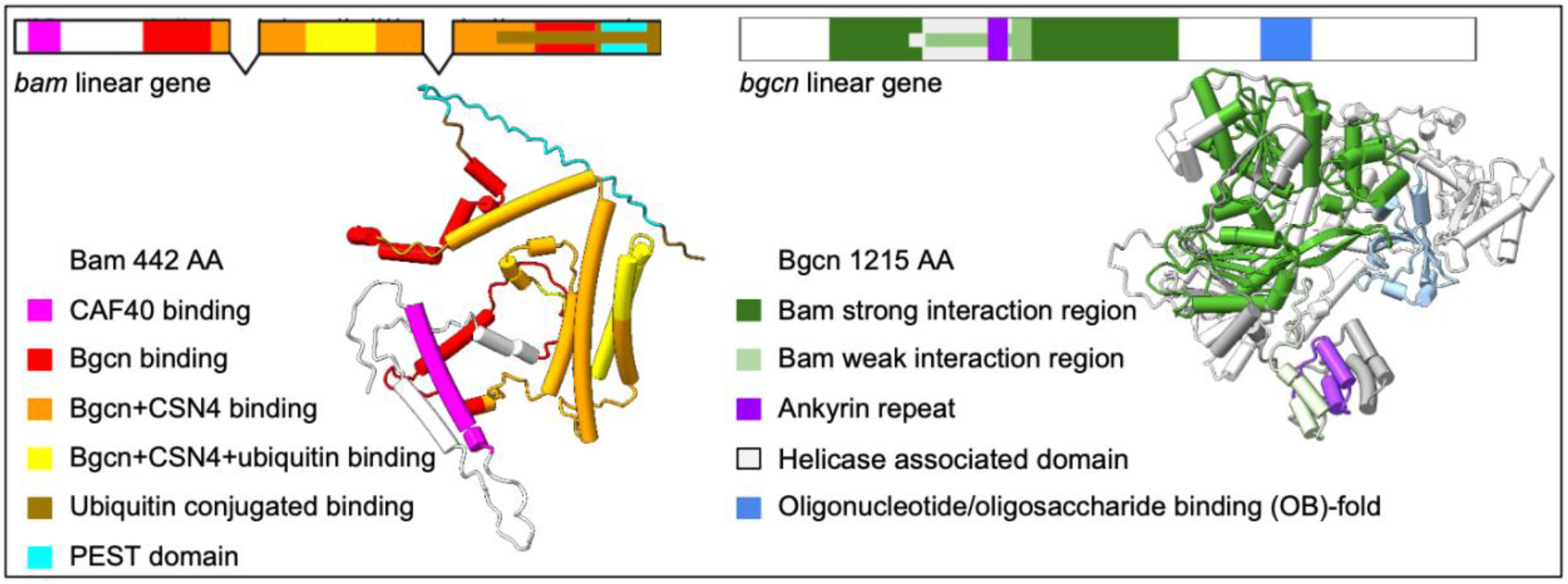
Predicted structures for *D. melanogaster* Bam and Bgcn with color-coded labels for their functional and binding regions

These studies have shown that *D. melanogaster* Bam has six identified functional regions (Fig. 1). The CAF40 binding motif (residues 13-36) (Ji et al. 2009; Sgromo et al. 2017), Bgcn binding region (residues 100-400) (Pan et al. 2014), Csn4 binding region (151-350) (Pan et al. 2014), free ubiquitin binding region (200-250) (Ji et al. 2017), conjugated ubiquitin binding region (300-442) (Ji et al. 2017), and a PEST domain (402-434) that marks the protein for rapid degradation (McKearin and Spradling 1990). *D. melanogaster* Bgcn has four biochemically identified regions, including one Bam binding region (Fig. 3). The Bam interaction region (residues 150-772), the helicase associated domain (300-442), the ankyrin repeat (407-439), and the oligonucleotide/oligosaccharide (OB) fold (862-945). The helicase associated domain present in Bgcn is a several-barrel structure found in RNA/DNA helicases (Xing et al. 2014) though insufficient alone for Bgcn to be a functional helicase (Ohlstein, 2000). The ankyrin-repeat sequence motif is thought to mediate protein-protein interactions and is critical for folding and stability (Li et al. 2006). The OB fold domain is a five-stranded β-barrel critical in protein-DNA and protein-protein interactions during DNA damage response (Flynn et al. 2010). The predicted structure for the Bam:Bgcn complex shows the more flexible and disordered Bam binds around the larger and more ordered Bgcn (Figure 4).

**Figure 4.**
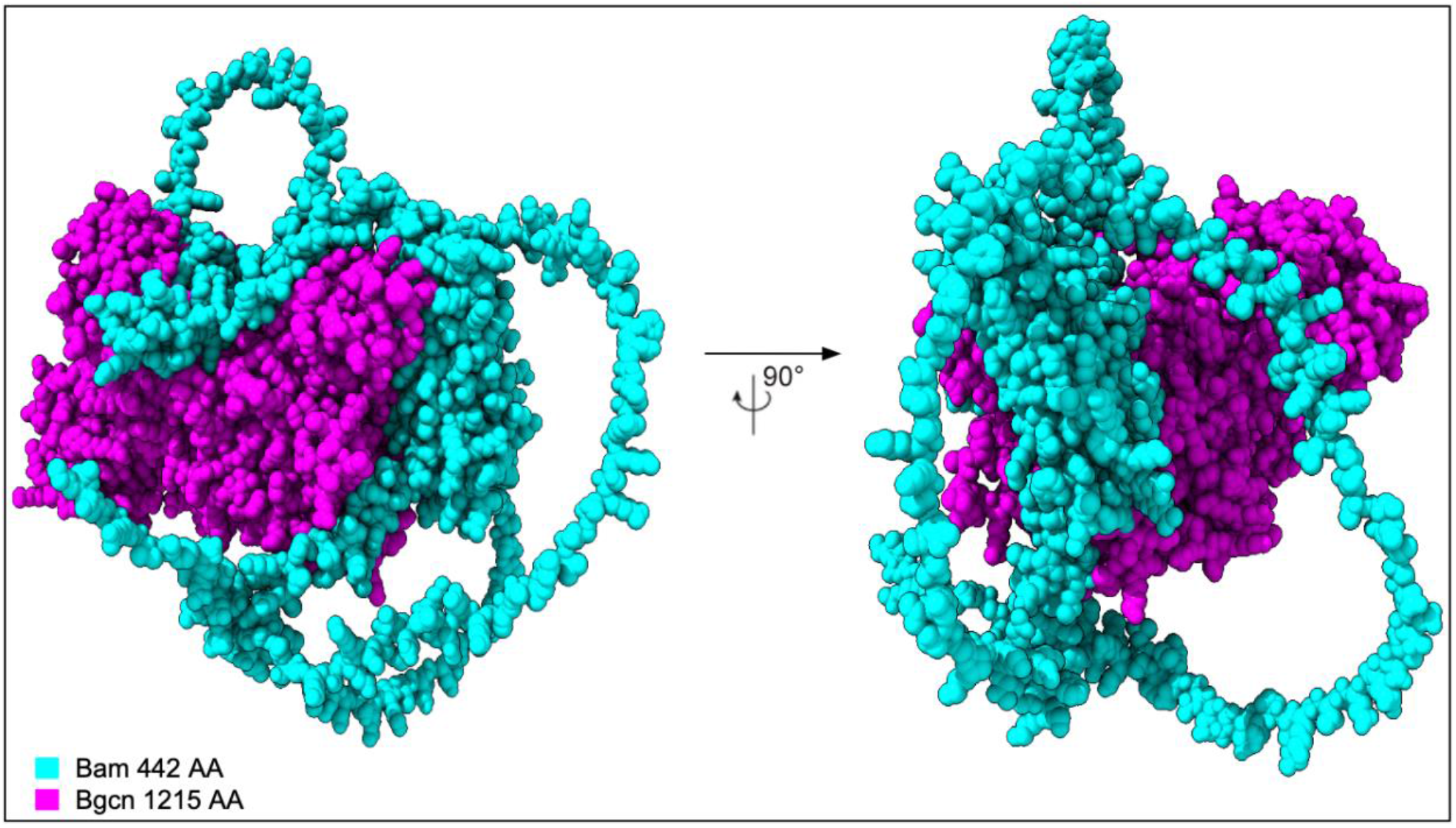
Space-filling predicted structure of the *D. melanogaster* Bam:Bgcn protein complex.

### Confidence in structural and positional predictions using plDDT, PAE, and ADOPT scores

To identify predicted structural changes in Bam and Bgcn proteins across the four species on which we focused, we examined paired predicted structural alignments of Bam-Bgcn complexes in each species. To effectively utilize the predicted protein structures, we used two measures of confidence: Predicted local distance test (plDDT) and predicted alignment error (PAE) (Guo et al. 2022). plDDT and PAE scores were used to specify which protein regions have high enough confidence scores to represent real structural predictions and ADOPT scores of structuredness were used to identify protein regions with high variance in structuredness between the species (S1).

plDDT scores reveal confidence in position of ∼60% amino acids of Bam and ∼95% of Bgcn. There are no significant differences (p <.05), using Fisher’s exact test, in confidence across species for either Bam or Bgcn alone as a percentage of the total protein or by functional region (Fig. 5, S1), and regions of confidence are largely consistent with Bam and Bgcn’s biochemically inferred functional regions. Regions of less confidence within the structures are predicted to be disordered or flexible linkages between ordered regions.

**Figure 5.**
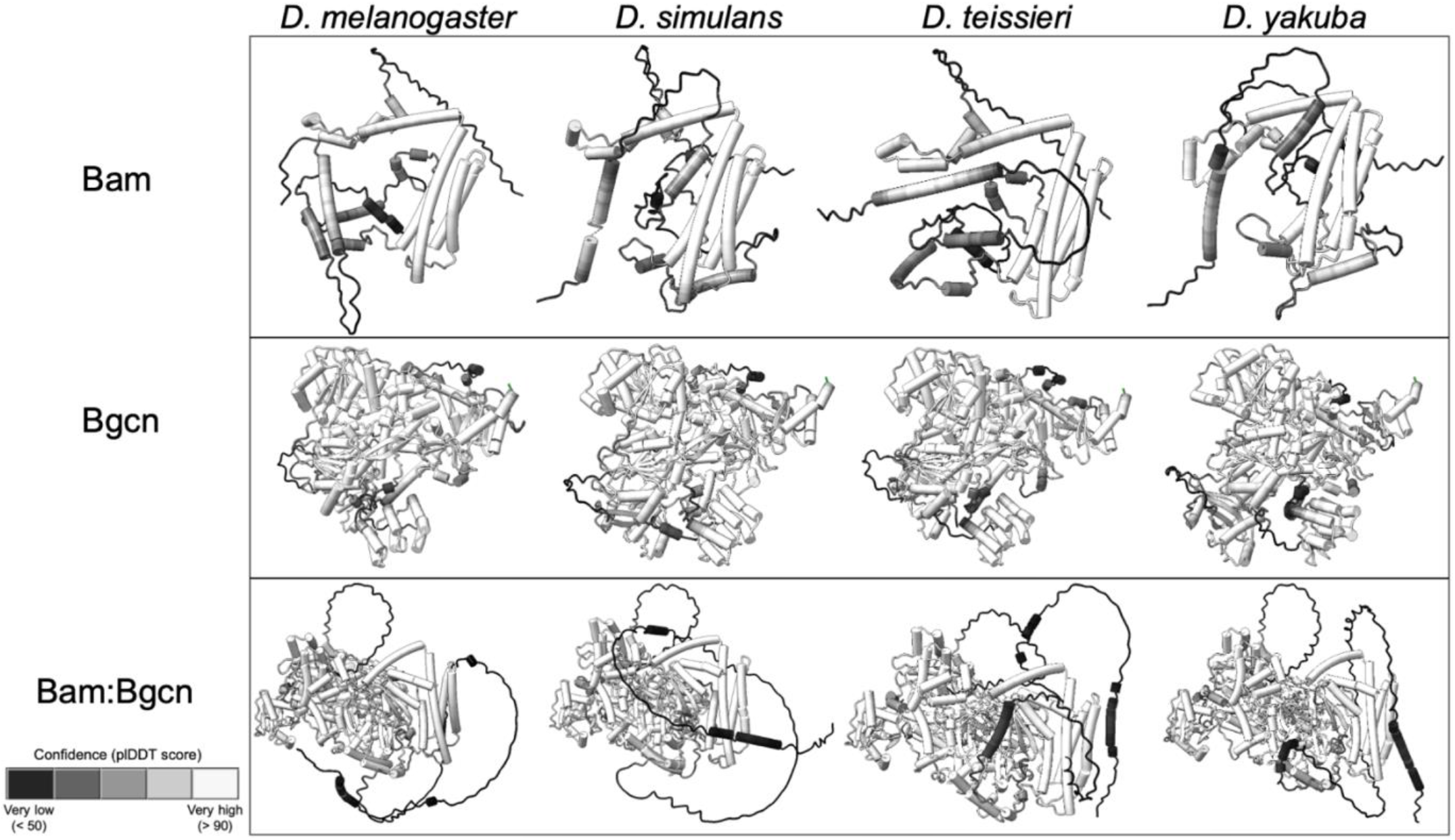
plDDT confidence scores for Bam, Bgcn, and Bam:Bgcn complex predicted structures for *D. melanogaster, D. simulans, D. teissieri*, and *D. yakuba*.

Bam:Bgcn protein complex plDDT scores also revealed no significant differences in regions for which AlphaFold was confident across all four species, for both Bam and Bgcn (Fig. 5, S1). There were differences in confidence between Bam and Bgcn alone vs in complex with one another. Specifically, there was less confidence in the N-terminal structure of Bam when it was bound to Bgcn, as compared to Bam alone, and there was overall greater confidence in the Bgcn structure when Bgcn was complexed with Bam.

PAE scores for Bam and Bgcn in each of the four species show low expected position errors between residues in confident, structured regions and high expected position errors between residues in disordered regions. This is also true for residues in Bam:Bgcn complexes (Fig. 6) which have low expected position error between residues in confident, structured regions within Bam and Bgcn as well as across Bam:Bgcn, particularly where the protein subunits are in close contact (S1, github.com/la424/bam_bgcn). The complex also has high expected position errors between the long, disordered regions of Bam and the structured regions of Bam as well as the Bgcn protein overall.

**Figure 6.**
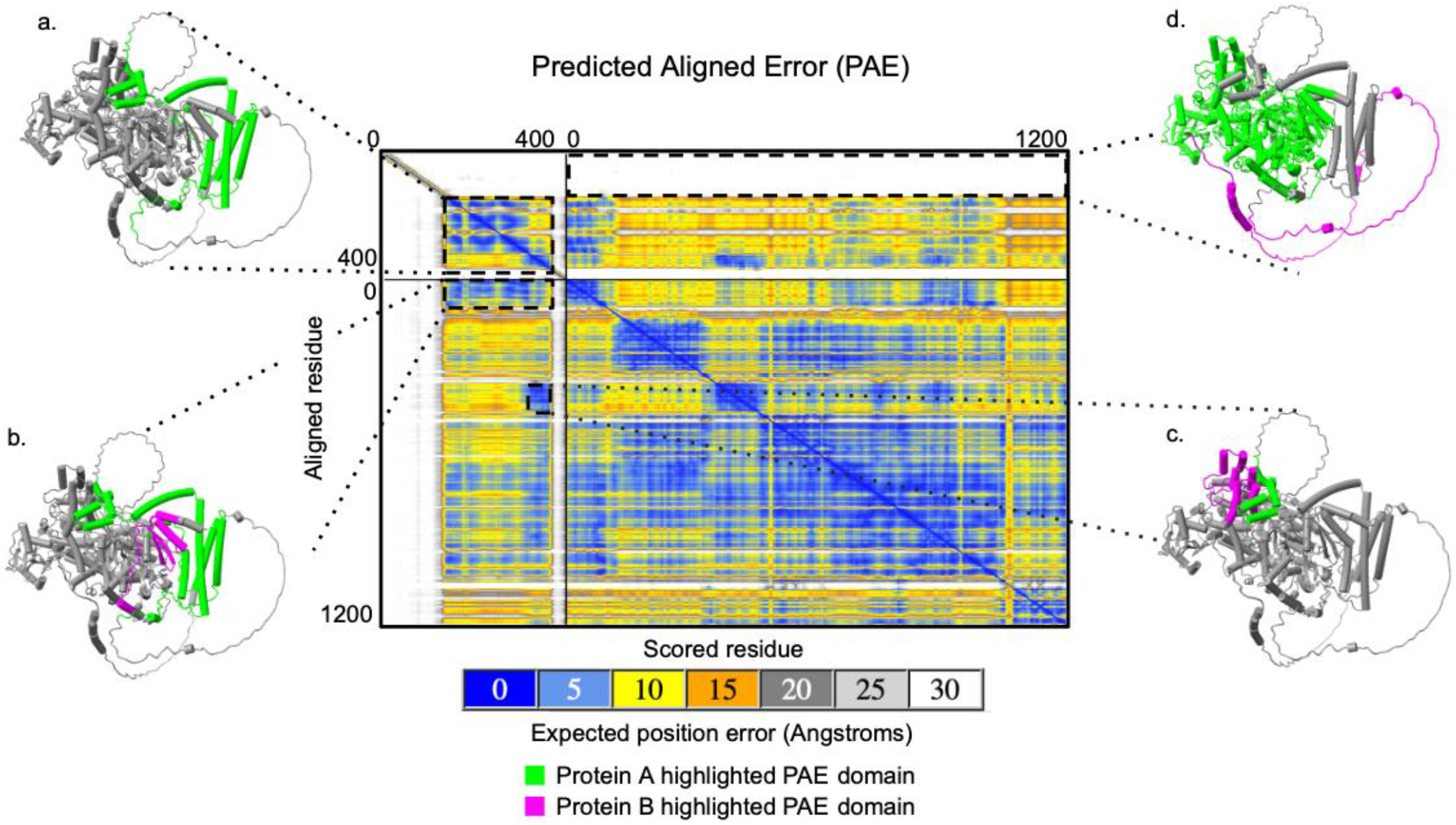
Predicted aligned error (PAE) plot for Bam:Bgcn complex residues in *D. melanogaster*. Lime green and magenta colors on the predicted structures are used to indicate whether the highlighted residues and PAE values are from one protein (Bam or Bgcn in lime green) or residues and PAE values across two proteins (Bam and Bgcn in lime green and magenta). a. Region with relatively low predicted error scores that highlight residues within Bam in which AlphaFold is confident in their relative positions. b and c. Regions with low predicted error scores that highlight residues from Bam and Bgcn in which AlphaFold is confident in their relative positions across proteins. d. Region with high predicted error scores that highlight residues from Bam and Bgcn in which AlphaFold is not confident in their relative positions across proteins.

Given the high proportion of Bam with predicted structural disorder by plDDT, we used an additional metric for predicting intrinsically disordered regions in Bam and Bgcn across the four Drosophila species. Attention based DisOrder PredicTor (ADOPT) is a machine learning based predictor of intrinsic structural disorder trained on curated datasets that originate from NMR chemical shift assignments of a mixture of structured and disordered proteins (Redl et al. 2023). The predicted Z-scores are residue-specific measures of structural order as observed in a database of more than 1400 intrinsically disordered proteins with known NMR (nuclear magnetic resonance) chemical shift assignments. NMR chemical shifts depend uniquely on the local environment and provide a probe, at single amino acid level, of the structure and dynamics of intrinsically disordered proteins. ADOPT structural predictions are more accurate than plDDT in disordered regions. Supplementing plDDT measured confidence with ADOPT scores helps develop a more comprehensive view of protein structure.

ADOPT scores for unbound Bam and Bgcn reveal no significant differences in predicted level of structuredness across included species (S1), and that level of structuredness tracks closely with confidence for Bam and Bgcn unbound assessed by plDDT.

### Evaluation of structural differences

The paired structural alignments were then evaluated for areas of predicted significant secondary structural and positional differences in confident regions of the protein structure. Bam, Bgcn, and Bam:Bgcn structures were directly superimposed using ChimeraX to identify residue-specific positional differences exceeding 2 Angstroms for the proteins and multimers between species (Meng et al. 2023). Predicted hydrogen bonds and AlphaFold contacts were also evaluated using ChimeraX. Focus was placed on whether there are structural changes across the species in general, or if there were species-specific structures, particularly in *D. teissieri*, the species whose functional data for Bam differ from the other three species’.

We used ChimeraX to compare structure between each species pair potentially identify structural differences between species. Paired structural alignments in ChimeraX for Bam unbound reveal no significant differences in predicted structure in confident regions across the four species. Paired structural alignment for Bam:Bgcn complex across the four species reveals no secondary structural differences and almost no positional differences in confident regions (Fig. 7, S1). There is one small *D. teissieri*-specific positional difference in a confident region in residues 344-348 in the four *Drosophila* species MSA (S1).

**Figure 7.**
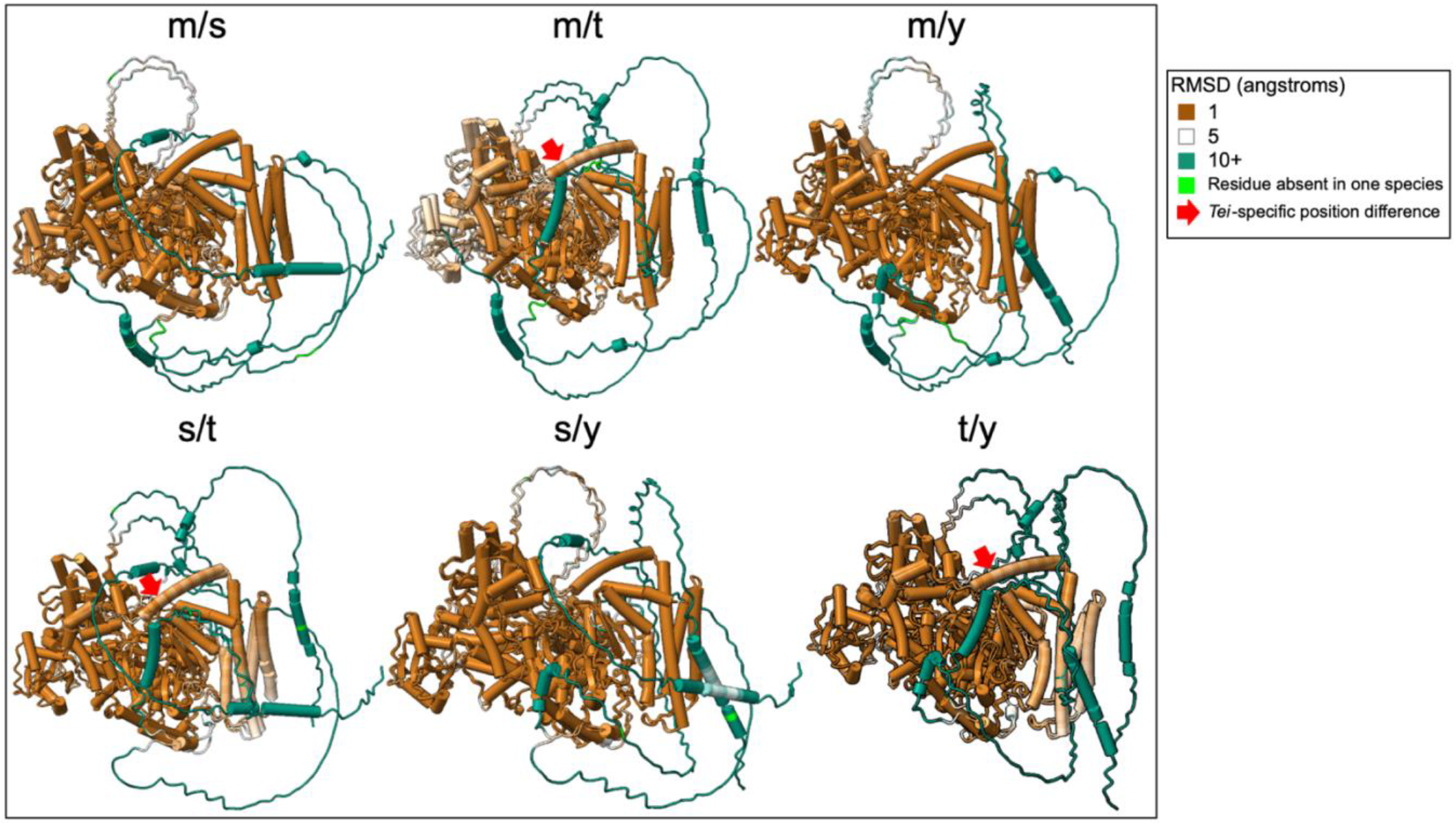
Paired structural alignments for Bam:Bgcn complex with *D. melanogaster, D. simulans, D. teissieri*, and *D. yakuba* indicated as m, s, t, and y respectively. The root mean square difference (RMSD) highlights the distance in angstroms between aligned species-specific residues. *D. teissieri*-specific residues with positional differences are highlighted with arrows.

Paired structural alignments for Bgcn unbound highlight predicted structural differences in confident regions between 325 and 475, but these differences are not species-specific. Paired alignments for Bgcn bound to Bam show a few 1-3 residue species-specific differences in confident regions for *D. teissieri* at positions 67, 889-890, 1011, 1118-1124, and 1151 in the four species MSA which are all predicted to be disordered linkages between ordered structures within Bgcn (S1).

For all four species we examined, predictive models of the Bam-Bgcn complex show Bam wrapping around the more structured Bgcn (Fig. 4), with Bam and Bgcn in contact in the biochemically identified (*D. melanogaster*) Bam and Bgcn binding regions (S1, S2, S3). Overall, there are no secondary structural changes across any of the predicted protein structures. There are some small predicted positional differences between species, but only one predicted difference in a confident, structured region in Bam bound at 344-348 for *D. teissieri*.

We used ChimeraX to identify predicted hydrogen bonds and AlphaFold contacts for Bam and Bgcn across the four species, and differences between species were compared using Fisher’s exact test (p<.05). There are no significant differences in number or distribution of predicted hydrogen bonding or AlphaFold contacts for Bam or Bgcn unbound or bound (S4, S5).

### Amino acid sequence differentiation among species

AlphaFold is an effective tool for predicting structure and position of ordered regions, but for disordered, low confidence regions, predicted structures are unreliable. *Bam* is a partially-disordered protein so these regions could contribute significantly to differences in *bam* function. To assess differences in disordered regions, appropriate predicted ancestral sequences were used to infer species-specific or lineage-specific amino acid substitutions. Amino acid changes from the three ancestral nodes were evaluated across species using several categories to search for potential patterns in significant differences. Differences were first evaluated and compared by total number of amino acid changes, by number in Bam and Bgcn functional and binding regions, by number in ordered and disordered regions, and by the ratio of amino acid changes in ordered regions to disordered regions. Fisher’s exact test followed by a Bonferroni correction for multiple hypothesis tests were then used to identify significant differences in amino acid changes for Bam and Bgcn across species. These substitutions were then evaluated based on volume and polarity using the Miyata scoring substitution matrix (Miyata et al. 1979). Scores range from 0, the most biochemically similar changes, to the least biochemically similar changes at 4. Miyata score differences were then evaluated for Bam across species using the same strategy (S3, S4, S5).

Bam pairwise amino acid sequence differences among the four *Drosophila* species included for predicted structure analysis (Fig 2b) range from 9% between *D. teissieri* and *D. yakuba* to 26% between *D. melanogaster* and *D. teissieri*. Also, there are more amino acid sequence differences between *D. melanogaster* and *D. simulans* at 16% than between *D. teissieri* and *D. yakuba* at 9%. This pattern holds when comparing the four included species to the ancestral sequence nodes, particularly node A as shown in the Bam amino acid linear alignment (Fig. 8.) Bam amino acid sequences for the four species differ from each other by up to 22% (*D. melanogaster* and *D. yakuba* / *D. teissieri*) and up to 33% from the Node A ancestral sequence (Fig 2b.) (Node A and *D. melanogaster* / *D. simulans*). There are more differences between the node A ancestral sequence and the *D. melanogaster* and *D. simulans* sequences (33% and 34% respectively) than the *D. yakuba* and *D. teissieri* sequences (22% and 19% respectively) (S1, github.com/la424/bam_bgcn). This variability is present throughout the length of the protein. Results for Bam predicted structures show Bam is predicted to have both disordered and structured regions in all four species examined (Fig 3). Most of the structured regions are included in the Bgcn, CSN4, and Ubiquitin binding regions of Bam, and these regions have higher sequence conservation across species. Two notable exceptions to the trend of high conservation in ordered regions are the residues 340-360 in the four species and node A multiple sequence alignment (msa) (github.com/la424/bam_bgcn)(residues 284-304 for just four species msa (S1)), a part of a long alpha helix in Bam’s Bgcn+CSN4 binding region, and residues 410-440 (four species + Node A MSA)(Github) (residues 353-383 (four species msa)), several small linked alpha helices in Bam’s Bgcn binding region (S1).

**Figure 8.**
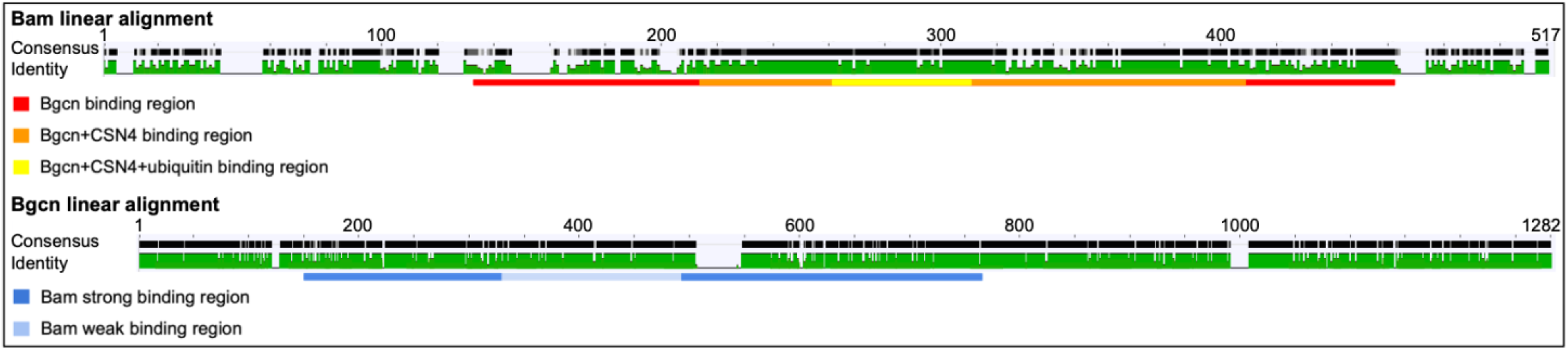
Bam and Bgcn linear amino acid sequence identity for *D. yakuba, D. teissieri, D. simulans*, and *D. melanogaster* aligned to the node A ancestral sequences with colored Bam and Bgcn binding regions. Level of per-residue conservation to the node A ancestral sequences is represented from no conservation as a light gray to total conservation in all included species as a thick black bar. Gaps are represented by a thin, gray line. The percent amino acid identity across species is represented below from red as the lowest % identity to green as 100% amino acid identity across included sequences with yellow bars of varying heights as intermediate values. Gaps are represented by a thin black line. Bam and Bgcn binding regions as determined in *D. melanogaster* are highlighted below linear alignments in colors specified in the key.

The Bgcn amino acid linear alignment (Fig 8) shows low levels of amino acid sequence variability across the four species that we examined throughout the protein, with slightly more variation in the Bam binding region. The Bgcn amino acid sequences differ from each other by up to 10% (*D. simulans* and *D. yakuba* / *D. teissieri*) and up to 13% from the Node A ancestral sequence (Fig 2) (Node A and *D. simulans*). Structures predicted for Bgcn (Fig. 3) show a largely structured protein with small, disordered linkages between regions of structure; this is seen in all four species we examined.

Bam amino acid sequences of the *four Drosophila* species were compared, and appropriate predicted ancestral sequences were used to infer species-specific or lineage-specific amino acid substitutions. For amino acid changes from ancestral sequence A, there are only significant differences between species across the *D. melanogaster* and *D. simulans* clade and the *D. yakuba* and *D. teissieri* clade for the total protein, functional regions, ordered regions, and disordered regions. These substitutions between select *Drosophila* species and ancestral sequence A were then evaluated based on biochemically similarity using the Miyata scoring substitution matrix (Miyata et al. 1979; referred to as Miyata score here) and compared across species. Amino acid changes with a Miyata score of 1 show a split pattern and amino acid changes with a Miyata score of 2 are significantly different between *D. melanogaster* and *D. teissieri*. Bam’s predicted Bgcn and Csn4 binding regions show significant differences between *D. simulans* and *D. teissieri* for Miyata score 1. There are no other significant differences in amino acid changes between the ancestral sequence A and the 4 species we examined, and no significant differences in amino acid sequence based on Miyata scores in any of the considered categories specific to the functionally distinct species *D. teissieri*. There are also no significant differences in amino acid changes from ancestral sequences B and C (S2, S3).

## DISCUSSION

Amino acid sequences for Bam, Bgcn, and predicted structures of Bam, Bgcn, and Bam bound to Bgcn were evaluated for significant differences across four *Drosophila* species using multiple sequence alignments, assessments of amino acid substitutions, structural alignments, hydrogen bonding, and AlphaFold contacts. Surprisingly, we found little significant predicted structural differences across the four species in regions of Bam or Bgcn where structures were predicted confidently despite the remarkable level of amino acid diversification between these four species. Bam:Bgcn complex predicted structures also showed points of contact between biochemically identified Bam and Bgcn binding regions giving more confidence in the structural predictions, particularly for regions of high structural and relative positional confidence based on plDDT and PAE scores. That said, there were abundant amino acid substitutions between the species, though none were specific to *D. teissieri*, the only species of the four in which Bam does not have a germline stem cell function. The predicted structures, hydrogen bonding, contact points were similar across all four species despite functional differences in *D. teissieri bam*, an amino acid substitution rate of up to 26% between the four species, and heterogenous signals of positive selection across *Drosophila*.

The Bam amino acid sequence shows more divergence than Bgcn across the *Drosophila* species that we examined. For most of the Bam and Bgcn sequences, we observe greater amino acid sequence divergence between species in regions of these proteins that are predicted to be structurally disordered. Two small structurally ordered regions in Bam however show elevated sequence divergence (Fig. 8.): residues 340-360 (four species + Node A MSA) (284-304 (four species MSA)) which are part of a long alpha helix in Bam’s Bgcn+CSN4 binding region, and residues 410-440 (four species + Node A MSA) (353-383 (four species MSA)) which form several small linked alpha helices in Bam’s Bgcn binding region. Whether there are functional consequences associated with changes in these regions is unknown. The 410-440 region of Bam does bind to Bgcn (Pan et al. 2014), so significant changes to the region could influence the binding between the two protein partners potentially impacting their function as a complex.

We did not find evidence that Bam structural changes underlie its altered function in *D. teissieri*. The lack of detectable structural change despite strong statistical signatures of positive selection and significant amino acid diversification across species raises the possibility that changes at *bam* among species are a mix of adaptive changes (possibly associated with some sort of arms race) coupled with compensatory amino acid substitutions necessary to retain a conserved 3-dimentional protein structure along the lines proposed in the Selection, Pleiotropy, and Compensation (SPC) model (Tokusicev and Wagner 2012). The SPC model for adaptive evolution predicts that adaptive change associated with one trait triggers deleterious pleiotropic effects in other traits or characteristics of function, and thus that additional selection occurs for compensatory substitutions that mitigate these deleterious pleiotropic effects (Baatz and Wagner 1997; Johnson and Porter 2001). An example of SPC is the evolution of drug resistance. Generally, drug resistance has a fitness cost if the drug is absent, but, when the drug is present, the general fitness cost is overshadowed by the benefits of resistance to the drug and resistant individuals survive. The secondary fitness costs of drug resistances are then compensated for by additional selection. The variability of *bam*’s role as the essential switch regulating a conserved developmental process (GSC differentiation and the production of gametes) across *Drosophila* species is also consistent with the concept of developmental systems drift (DSD) which is the change in the developmental mechanics of a phenotypically conserved trait (True and Haag 2001; Haag 2007).

The signals of positive selection at *bam* do not necessarily indicate that selection is changing *bam*’s role as GSC differentiation switch. Since *bam* plays several key roles including in gametogenesis (McKearin and Spradling 1990), hematopoiesis (Tokusumi et al. 2011), and gut homeostasis (Ji et al. 2019), there could be multiple candidate primary drivers of the positive selection observed at *bam* in numerous species lineages. Evasion of reproductive manipulation of gametogenesis by the endosymbiont *Wolbachia* remains a strong possibility as a primary driver (Flores et al. 2015, Russell et al. 2023). Secondary, compensatory changes at *bam* mitigating potential deleterious pleiotropic effects on structure and/or other functions of Bam are also possible (Werren et al. 2008; Kaur et al. 2021; Wenzel and Aquadro 2023). The lack of predicted structural differences across species for Bam and its complex with Bgcn is consistent with the view that changes in *bam*’s role in GSC differentiation are not driving amino acid diversification. The predicted structural conservation raises the possibility that at least some amino acid changes could be compensatory changes to conserve the protein and protein complex secondary structures. The positions of amino acid changes between species also are also consistent with the conservation of Bam:Bgcn binding in all 4 species. Detection of potentially compensatory changes at Bam:Bgcn binding sites even within *D. teissieri*, in which Bam is not essential for GSC differentiation (Bubnell et al. 2022) could be because of an alternative role that Bam:Bgcn binding executes beyond the GSC switch function or because most of these compensatory changes occurred before *bam* became no longer essential for switching GSC differentiation in *D. teissieri*.

The combination of adaptive and compensatory change in the SPC model of course complicate detection of the source of primary adaptive selection (Pavlicev and Wagner 2012). Amino acid substitutions consistent with a signature of adaptive evolution could actually be compensatory changes instead of a state change in the trait. More and more genes are being identified as pleiotropic with recent estimates putting the average degree of pleiotropy ranging up to 21 traits per locus (Wagner and Zhang 2011). In the SPC model, this suggests that secondary compensatory changes could be up to 21 times more likely in the genome than adaptive changes representing a change in trait state (Pavlicev and Wagner 2012). In order to sort detected signals of positive selection for state changes versus compensatory changes, the accumulation of many more *bam* (and *bgcn*) sequences from diverse species and lineages will be needed. Together with results from our comparative predicted structural analysis, the use of computational and phylogenetic tools to distinguish primary from compensatory amino acid changes may be possible (e.g., Dutheil and Galtier 2007; Chaurasia and Duthiel 2022).

## Acknowledgements

We thank Mariana Wolfner for thoughtful comments on an earlier version of the manuscript. This work was supported by National Institute of Health (United States) R01-GM095793 to Charles F. Aquadro.

## References

Baatz M and Wagner G. 1997. Adaptive inertia caused by hidden pleiotropic effects. Theor. Popul. Biol. 51:49–66. 10.1006/tpbi.1997.1294

Bauer DuMont VL, Flores HA, Wright MH, Aquadro CF. 2007. Recurrent positive selection at Bgcn, a key determinant of germ line differentiation, does not appear to be driven by simple coevolution with its partner protein bam. Mol Biol Evol. 24(1):182–191. 10.1093/molbev/msl141

Bubnell JE, Ulbing C, Begne PF, Aquadro CF. 2022. Functional divergence of the bag-of-marbles gene in the Drosophila melanogaster species group. Molecular Biology and Evolution. 39(7):msac137. 10.1093/molbev/msac137

Camacho, C., Coulouris, G., Avagyan, V., Ma, N., Papadopoulos, J., Bealer, K., and Madden, T.L. 2009. BLAST+: architecture and applications. BMC Bioinformatics, 10, 421. 10.1186/1471-2105-10-421

Chaurasia S and Duthiel J. 2022. The structural determinants of intra-protein compensatory substitutions. Molecular Biology and Evolution 39 Apr 11;39(4):msac063. 10.1093/molbev/msac063

Chau J, Kulnane LS, Salz HK. 2012. Sex-lethal enables germline stem cell differentiation by down-regulating Nanos protein levels during Drosophila oogenesis. Proc Natl Acad Sci USA.;109:9465–9470. 10.1073/pnas.1120473109.

Chen D, McKearin D. 2003. Dpp signaling silences bam transcription directly to establish asymmetric divisions of germline stem cells. Curr Biol. 13(20):1786– 1791. 10.1016/j.cub.2003.09.033.

Chen D, Wu C, Zhao S, Geng Q, Gao Y, Li X, Zhang Y, Wang Z. 2014. Three RNA binding proteins form a complex to promote differentiation of germline stem cell lineage in Drosophila. PLoS Genet. 10(11):e1004797. 10.1371/journal.pgen.1004797.

Civetta A, Rajakumar SA, Brouwers B, Bacik JP. 2006. Rapid evolution and gene-specific patterns of selection for three genes of spermatogenesis in Drosophila. Mol Biol Evol. 23(3):655–662. 10.1093/molbev/msj074.

Choi JY, Aquadro CF. 2014. The coevolutionary period of Wolbachia pipientis infecting Drosophila ananassae and its impact on the evolution of the host germline stem cell regulating genes. Mol Biol Evol. 31(9):2457–2471. 10.1093/molbev/msu204.

Duthiel J and Galtier N. 2007. Detecting groups of coevolving positions in a molecule: a clustering approach. BMC Evolutionary Biology 7:242. 10.1186/1471-2148-7-242.

Edgar R. 2004. MUSCLE: multiple sequence alignment with high accuracy and high throughput. Nucleic Acid Res. 32(5):1792–1797. 10.1093/nar/gkh340.

Evans, R. et al. 2021. “Protein complex prediction with AlphaFold-Multimer.” bioRxiv. 10.1101/2021.10.04.463034.

Flores HA, Bubnell JE, Aquadro CF, Barbash DA. 2015. The Drosophila bag of marbles gene interacts genetically with Wolbachia and shows female-specific effects of divergence. PLoS Genet. 11:e1005453. 10.1371/journal.pgen.1005453.

Flores HA, DuMont VLB, Fatoo A, Hubbard D, Hijji M, Barbash DA, Aquadro CF. 2015. Adaptive evolution of genes involved in the regulation of germline stem cells in Drosophila melanogaster and D. simulans. G3 Genes, Genomes, Genet. 5(4):583– 592. 10.1534/g3.114.015875.

Flynn R. et al. 2010. Oligonucleotide/Oligosaccharide-Binding (OB) fold proteins: A growing family of genome guardians. Crit Rev Biochem Mol Biol. Aug; 45(4):266–275. 10.3109/10409238.2010.488216.

Freilich S. et al. 1999. The COP9 signalosome is essential for development of Drosophila melanogaster. Current Biology. Oct 21;9(20):1187-90. 10.1016/S0960-9822(00)80023-8.

Fu Z, Geng C, Wang H, Yang Z, Wenig C, Li H, et al. 2015. Twin promotes the maintenance and differentiation of germline stem cell lineage through modulation of multiple pathways. Cell Rep. 13:1366–1379. 10.1016/j.celrep.2015.10.017.

Gleason RJ, Anand A, Kai T, Chen X. 2018. Protecting and diversifying the germline. Genetics 208(2):435–471. 10.1534/genetics.117.300208.

Guo, H.-B. et al. 2022. AlphaFold2 models indicate that protein sequence determines both structure and dynamics. Scientific Reports, 12(1):10696. 10.1038/s41598-022-14382-9.

Haag ES. 2007. Compensatory vs. pseudocompensatory evolution in molecular and developmental interactions. Genetica. 129:45–55. 10.1007/s10709-006-0032-3.

Insco ML, Leon A, Tam CH, McKearin DM, Fuller MT. 2009. Accumulation of a differentiation regulator specifies transit amplifying division number in an adult stem cell lineage. Proc Natl Acad Sci. 106(52):22311–22316. 10.1073/pnas.0912454106.

Ji S, Li C, Hu L, Liu K, Mei J, Luo Y, et al. 2017. Bam-dependent deubiquitinase complex can disrupt germ-line stem cell maintenance by targeting cyclin A. Proc Natl Acad Sci USA 114:6316–6321. 10.1073/pnas.1619188114.

Ji, S., Luo, Y., Cai, Q., Cao, Z., Zhao, Y., Mei, J., Li, C., Xia, P., Xie, Z., Xia, Z., Zhang, J., Sun, Q., Chen, D., 2019. LC domain-mediated coalescence is essential for Otu enzymatic activity to extend drosophila lifespan. Molular Cell 74:363–377. 10.1016/j.molcel.2019.02.004

Johnson NA and Porter AH. 2001. Toward a new synthesis: population genetics and evolutionary developmental biology. Genetica, 112-113: 45–58.

Jumper, J. et al. 2021. “Highly accurate protein structure prediction with AlphaFold.” Nature, 596, pages 583–589. 10.1038/s41586-021-03819-2.

Kahney EW, Snedeker JC, Chen X. 2019. Regulation of Drosophila germline stem cells. Curr Opin Cell Biol. 60:27–35. 10.1016/j.ceb.2019.03.008.

Kaur R, Shropshire JK, Cross KL, Leigh B, Mansueto AJ, Steward V, et al. 2021. Living in the endosymbiotic world of Wolbachia: A centennial review. Cell Host Microbe 29(6):879–93. 10.1016/j.chom.2021.03.006.

Li J. et al. 2006. Ankyrin repeat: a unique motif mediating protein-protein interactions. Biochemistry. 45(51):15168–78. 10.1021/bi062188q.

Li Y, Minor NT, Park JK, McKearin DM, Maines JZ. 2009. Bam and Bgcn antagonize Nanos-dependent germ-line stem cell maintenance. Proc Natl Acad Sci USA. 106(23):9304–9309. 10.1073/pnas.0901452106.

Malik S, Jang W, Kim JY, Kim C. 2020. Mechanisms ensuring robust repression of the Drosophila female germline stem cell maintenance factor Nanos via posttranscriptional regulation. The FASEB Journal 34:11421–11430. 10.1096/fj.202000656R.

McKearin DM, Spradling AC. 1990. Bag-of-marbles: A Drosophila gene required to initiate both male and female gametogenesis. Genes Dev. 4(12 B):2242–2251. 10.1101/gad.4.12b.2242.

Meng EC, Goddard TD, Pettersen EF, Couch GS, Pearson ZJ, Morris JH, Ferrin TE. 2023. UCSF ChimeraX: Tools for structure building and analysis. Protein Sci. 32(11):e4792. 10.1002/pro.4792.

Mills JE and Deam PM. 1996. Three-dimensional hydrogen-bond geometry and probability information from a crystal survey. Journal of Computer-Aided Molecular Design 10:607–622. 10.1007/BF00134183.

Mirdita M. et al. 2022. “ColabFold: Making protein folding accessible to all.” Nature Methods, 19:679–682. 10.1038/s41592-022-01488-1.

Miyata T. et al. 1979. Two types of amino acid substitutions in protein evolution. Journal of Molecular Evolution. 12:219–236. 10.1007/BF01732340.

O’Leary NA et al. 2016. Reference sequence (RefSeq) database at NCBI: current status, taxonomic expansion, and functional annotation. Nucleic Acids Res. 44(D1):D733-45. 10.1093/nar/gkv1189.

Ohlstein B. et al. 2000. The Drosophila cystoblast differentiation factor, benign gonial cell neoplasm, is related to DExH-box proteins and interacts genetically with bag-of-marbles. Genetics. 155(4):1809–19. 10.1093/genetics/155.4.1809.

Pan L, Wang S, Lu T, Weng C, Song X, Park JK, et al. 2014. Protein competition switches the function of COP9 from self-renewal to differentiation. Nature 514:233– 236. 10.1038/nature13562.

Pavlicev M and Wagner G. 2012. A model of developmental evolution: selection, pleiotropy, and compensation. Trends in Ecology and Evolution Volume 27, Issue 6, June. 10.1016/j.tree.2012.01.016.

Redl, Istvan, et al. 2023. ADOPT: intrinsic protein disorder prediction through deep bidirectional transformers. Nucleic Acids Res. 1;5(2):qad041. 10.1093/nargab/lqad041.

Russell, SL, Castillo, JR and Sullivan, WT. 2023. Wolbachia endosymbionts manipulate the self-renewal and differentiation of germline stem cells to reinforce fertility of their fruit fly host. PLOS Biology 21(10):e3002335. 10.1371/journal.pbio.3002335

Sgromo A, Raisch T, Bawankar P, Bhandari D, Chen Y, Kuzuoğlu-Öztürk D, et al. 2017. A CAF40-binding motif facilitates recruitment of the CCR4-NOT complex to mRNAs targeted by Drosophila Roquin. Nat Commun. 8:14307. 10.1038/ncomms14307.

Sgromo A. et al. 2018. Drosophila Bag-of-marbles directly interacts with the CAF40 subunit of the CCR4-NOT complex to elicit repression of mRNA targets. RNA. 24(3):381–395. 10.1261/rna.064584.117.

Tokusumi T. et al. 2011. Germ line differentiation factor Bag of Marbles is a regulator of hematopoietic progenitor maintenance during Drosophila hematopoiesis. Development. 138(18):3879–84. 10.1242/dev.069336.

True JR and Haag ES. 2001. Developmental system drift and flexibility in evolutionary trajectories. Evol. Dev. 3:109–119. 10.1046/j.1525-142x.2001.003002109.x.

Wagner GP and Zhang J. 2011. The pleiotropic structure of the genotype–phenotype map: the evolvability of complex organisms. Nat. Rev. Genet. 12:204–213. 10.1038/nrg2949.

Weeratunga, S. et al. 2023. Interrogation and validation of the interactome of neuronal Munc18-interacting Mint proteins with AlphaFold2. J Biol Chem. 300(1):105541. 10.1016/j.jbc.2023.105541.

Wenzel M and Aquadro F. 2023. Wolbachia infection at least partially rescues the fertility and ovary defects of several new Drosophila melanogaster bag of marbles protein-coding mutants. PLOS Genetics 8:23. 10.1101/2023.03.20.532813.

Werren JH, Baldo L, Clark ME. 2008. Wolbachia: master manipulators of invertebrate biology. Nat Rev Microbiol. 6:741–751. 10.1038/nrmicro1969.

Xie T. 2013. Control of germline stem cell self-renewal and differentiation in the Drosophila ovary: Concerted actions of niche signals and intrinsic factors. Wiley Interdiscip Rev Dev Biol. 2(2):261–273. 10.1002/wdev.60.

Xing L. et al. 2014. Helicase associated 2 domain is essential for helicase activity of RNA helicase A. BBA Proteins and Proteomics. Volume 1844, Issue 10, 1757–1764. 10.1016/j.bbapap.2014.07.001.

Xu, J., & Zhang, Y. 2010. How significant is a protein structure similarity with TM-score = 0.5? Bioinformatics, 26(7), 889–895. 10.1093/bioinformatics/btq066.

